# High salinity activates CEF and attenuates state transitions in both psychrophilic and mesophilic *Chlamydomonas* species

**DOI:** 10.1101/2022.03.14.484132

**Authors:** Isha Kalra, Xin Wang, Ru Zhang, Rachael Morgan-Kiss

## Abstract

In the last decade, studies have revealed the importance of PSI-driven cyclic electron flow (CEF) in stress acclimation in model organisms like *C. reinhardtii*; however, these studies focused on transient, short-term stress. In addition, PSI-supercomplexes are associated with CEF during state transition response to short-term stress. On the other hand, the role of CEF during long-term stress acclimation is still largely unknown. In this study, we elucidate the involvement of CEF in acclimation response to long-term high salinity in three different *Chlamydomonas* species displaying varying salinity tolerance. We compared CEF rates, capacity for state transitions, and formation of supercomplexes after salinity acclimation in the model mesophile *C. reinhardtii* and two psychrophilic green algae *C. priscuii* (UWO241) and *C. sp*. ICE-MDV. CEF was activated under high salt in all three species, with the psychrophilic *Chlamydomonas* spp. exhibiting the highest CEF rates. High salt acclimation was also correlated with reduced state transition capacity and a PSI-supercomplex was associated with high CEF. We propose that under long-term stress, CEF is constitutively activated through assembly of a stable PSI-supercomplex. The proteomic composition of the long-term PSI-supercomplex is distinct from the supercomplex formed during state transitions, and its presence attenuates the state transition response.

## INTRODUCTION

Salinity is a challenging abiotic stress encountered by plants and algae. With intensified agricultural practices, increased soil salinization has caused significant losses in crops and global productivity (Morton et al., 2019; Welle and Mauter, 2017). Excess salinity causes ion toxicity and disturbs osmotic balance (Hasegawa et al., 2000; Kumar et al., 2018). Additionally, under high salt conditions, photosynthetic organisms can experience loss of membrane organization, inhibition of photosynthesis, increased reactive oxygen species, and disruptions in nutrient acquisition. Responses to high salinity include: (i) changes in cell wall turgor or volume, ii) selective uptake of ions by organelles such as vacuoles, and (iii) active exclusion of sodium ions and/or accumulation of compatible osmolytes such as glycerol in the cell to maintain ion homeostasis (Goyal, 2007; He et al., 2015, Figler et al. 2019).

Exposure to any abiotic stress leads to disruptions in cell homeostasis, including over-reduction of the photosynthetic electron transport chain (PETC). This over-reduction of PETC can lead to production of ROS or photoinhibition, and organisms must balance this redox imbalance to maintain growth and photosynthesis (Hüner et al., 2012). In their natural environments, organisms encounter a myriad of environmental stresses (e.g. high light, low temperatures, high salinity, nutrient deficiency), which may last for a few minutes (short-term or transient) or persist for days to years (long-term) (Kono & Terashima, 2014). Photosynthetic organisms respond to these environmental perturbations either through short-term acclimatory responses such as state-transitions or long-term reorganization of the photosynthetic apparatus or shifts in downstream carbon metabolism. PSI-driven cyclic electron flow (CEF), is essential for balancing energy needs for carbon fixation and in survival under stress by supplying additional ATP and/or activating photoprotection, thus also balancing the redox status (Kramer & Evans, 2011; Suorsa, 2015). Although our knowledge CEF has grown rapidly in the past decade, the current understanding of the role of CEF in stress acclimation has been mainly restricted to treatments over short time scales. There is an underappreciation for CEF mechanism and function during long-term stress acclimation or adaptation to permanently stressful environments (DalCorso et al., 2008; Iwai et al., 2010; Lucker & Kramer, 2013; Takahashi et al., 2013; Takahashi et al., 2016).

While the exact mechanism of CEF initiation is still debated, formation of thylakoid protein supercomplexes appear to play an important role in the activation of CEF (Minagawa, 2016). In the last decade, the contribution of PSI-supercomplexes in initiating CEF has been extensively studied in model organisms such as *C. reinhardtii* (Alric, 2010; Iwai et al., 2010; Minagawa, 2016; Steinbeck et al., 2018; Terashima et al., 2012). State transition conditions (ie. short-term exposure to a condition causing over-reduction of the PETC) have been implicated in assembly of the supercomplex and activation of transiently high CEF (Iwai et al., 2010; Minagawa, 2016). The supercomplex of *C. reinhardtii* cells in State 2 was shown to be composed of PSI-LHCI, LHCII, cytochrome b_6_f (cyt b_6_f), FNR (ferredoxin NADP reductase), PGRL1 (proton gradient like protein 1), CAS (calcium sensing protein) and other minor proteins (Z. Huang et al., 2021; Steinbeck et al., 2018; Takahashi et al., 2016; Terashima et al., 2012).

Many processes involved in salinity response require additional ATP; therefore, organisms exposed to high salt (both halophyte and non-halophyte) deal with continuous pressure to deal with high energy demands. In addition to playing a key role in photoprotection, CEF can also provide extra ATP. CEF constitutes a major pathway by which photosynthetic organisms fine tune the ATP/NADPH ratio based on downstream energetic demands (Suorsa, 2015). A significant body of research has been focused on salinity stress response in non-halophyte models, such as *C. reinhardtii.* (Neelam & Subramanyam, 2013; Sudhir & Murthy, 2004; Wang et al., 2018; Perrineau et al., 2014; Sithtisarn et al., 2017). Much of the work on salinity stress in *C. reinhardtii* was conducted under mixotrophic growth conditions, which confounds understanding how the PETC responds to salinity due to bypassing of photosynthetic growth in presence of acetate (Heifetz et al., 2000). In addition, halophilic and halotolerant algae (eg: *Dunaliella salina*) and plants (eg: *Atriplex, Mesembrayanthemum crystallinum*) can provide additional insights regarding the full potential of photosynthetic organisms to maintain rapid growth at high salinities (Greenway & Munns, 1980). *D. salina*, a dominant primary producer in hypersaline environments (Oren, 2014), is one of the few photosynthetic algal models for high salinity adaptation (Cowan et al., 1992). Direct comparisons between the reference alga *C. reinhardtii* and stress-tolerance organisms such as the halophyte *D. salina* are problematic due to their distant relatedness. There is a growing appreciation for efforts to expand studies to focus on a repertoire of ‘wild’ *Chlamydomonas* spp. exhibiting broad tolerances to environmental disturbances, including low temperatures, extremes in pH, nutrient-poor habitats, and desiccation (Grossman, 2021).

Non-model organisms that are adapted to very long-term stress lasting 100s-1000s of years represent under-exploited reservoirs of novel adaptive mechanisms (Dolhi et al., 2013). One such organism is the Antarctic psychrophile *Chlamydomonas priscuii* UWO241 (UWO241 henceforth), which was isolated from deep photic zone of the ice-covered lake Bonney in the McMurdo Dry Valleys (Morgan et al., 1998; Stahl-Rommel et al., 2021). Extensive studies on this organism have demonstrated that it is a psychrophilic, halotolerant green alga with a reorganized photosynthetic apparatus that includes PSII with a large light harvesting antenna, matched by PSI complexes with diminished antenna size (Dolhi et al., 2013; Morgan-Kiss et al., 2002b; Morgan et al., 1998; Szyszka et al., 2007). Some short-term acclimation strategies, including state transitions, appear to have been lost during adaptation to permanent and stable extreme stress (Morgan-Kiss et al., 2002a). As a consequence of adaptation to a native environment of hypersalinity and permanent cold, UWO241 exhibits sustained high rates of CEF, which is associated with assembly of a stable PSI-supercomplex (Morgan-Kiss et al., 2002; Morgan et al., 1998; Szyszka et al., 2007). Szyszka *et. al* (2015) detected proteins of PSI, cyt b_6_f, as well as two novel phosphor-proteins, FtsH and a PsbP-like protein in the UWO241 supercomplex (Szyszka-Mroz et al., 2015). Later, Kalra et al. (2020) improved the yield of the UWO241 PSI-supercomplex and reported the presence of several additional polypeptides, including several PSI subunits, chloroplastic ATP synthase, CAS and PGRL1. The constitutively high CEF appears to provide multiple advantages to UWO241, including extra ATP and a strong capacity for NPQ (Kalra et al., 2020).

More recently, a second photopsychrophile, named *Chlamydomonas* sp. ICE-MDV (ICE-MDV henceforth) was isolated from the Antarctic lake Bonney. Unlike UWO241, ICE-MDV resides in the shallow photic zone of the water column where it receives high light, but reduced salinity and very low nutrients (Li & Morgan-Kiss, 2019; Li et al., 2016). ICE-MDV is also psychrophilic; however, it exhibits some distinct physiological differences from UWO241 in its photosynthetic apparatus. For example, unlike UWO241, whole cells of ICE-MDV exhibit detectable levels of PSI low temperature fluorescence (Cook et al., 2019).

Previous studies have shown that adaptation to high salinity in the psychrophile UWO241 is manifested as formation of a novel PSI-supercomplex to support sustained high rates of CEF. We thus hypothesize that CEF and supercomplexes are important in long-term salinity stress acclimation. Here we investigated whether increased CEF may constitute a common strategy to fine tune redox status of PETC and provide extra ATP under long-term salinity stress in mesophilic and other psychrophilic *Chlamydomonas* species. We then hypothesized that acclimation to long-term salinity stress inhibits the capacity for state transitions through constitutive upregulation of PSI-supercomplex associated CEF. To test these hypotheses, we compared CEF rates, state transition ability, and looked for the presence of supercomplexes after long-term salinity acclimation in the salt sensitive reference strain *C. reinhardtii* as well as the two Antarctic algae adapted to moderate (ICE-MDV) and very high (UWO241) salinity in their native habitat.

## METHODS

### Culture conditions, growth physiology

Three different *Chlamydomonas* species were used in this study: *C*. *priscuii* (UWO241; strain CCMP1619), *C*. sp. ICE-MDV (ICE-MDV) and the model *C. reinhardtii* (strain UTEX 90). All three species were first grown in Bold’s Basal Media (BBM, 0.43 mM NaCl) (Low salt, LS). Based on previous studies, UWO241 and ICE-MDV cultures were grown under a temperature/irradiance regime of 8°C/50 μmol photons m^−2^ s^−1^ (Cook et al., 2019; Morgan et al., 1998). *C.reinhardtii* UTEX 90 was grown in BBM (LS) at 20°C/100 μmol photons m^−2^ s^−1^. All cultures were grown in 250 ml glass pyrex tubes in temperature regulated aquaria under a 24 h light cycle and were continuously aerated with sterile air supplied by aquarium pumps (Morgan-Kiss et al., 2002).

For the salinity tolerance study, cultures were grown in increasing concentration of NaCl supplemented BBM (0.43-700 mM NaCl for UWO241 and ICE-MDV, 0-200 mM NaCl for *C. reinhardtii*). Growth was monitored by optical density at wavelength of 750 nm. Maximum growth rates were calculated using natural log transformation of the optical density values during the exponential phase. Three biological replicates were performed.

For salinity stress acclimation, cultures were grown in maximum tolerated salinity levels and sub-cultured after reaching log-phase, at least 2-3 times. All subsequent experiments were conducted on low salinity (LS) and high salinity acclimated (HS) log-phase cultures.

### Room-temperature PSII chlorophyll fluorescence measurements

Photosynthetic measurements were conducted using room temperature PSII chlorophyll fluorescence through Dual PAM-100 instrument (Walz, Germany). Briefly, 2 ml of exponentially grown cultures were dark adapted using far-red light for 2 min prior to the measurement. For steady-state analysis, we used induction curves to measure maximum capacity of photosynthesis (F_v_/F_m_), photosynthetic yield (YPSII), non-photochemical quenching (YNPQ) and photochemical yield (qP) with actinic light set at growth light intensities for each species. Light curves in Dual-PAM were also conducted to measure the change in capacity of NPQ with increasing light levels.

### State transition induction

State transition experiments were conducted on both low and high salinity acclimated cultures. Briefly, cultures were harvested in the mid-log phase and induced in either state 1 or state 2 through addition of chemical inhibitors as described before (Iwai et al., 2010). For state 1 induction, mid-log phase cells were incubated in 10 µM DCMU to completely oxidize the PQ pool prior to measurement. For state 2 induction, cells were incubated in 5 µM FCCP for 20 min. State transition response was measured through either 77 K spectra or PSII fluorescence as described below.

### Low temperature (77K) fluorescence spectra

Low temperature (77K) fluorescence spectra were conducted as previously described (Morgan et al. 1998). Briefly, log-phase cultures or isolated Chl-protein complexes were dark adapted for 10 mins and flash frozen in liquid N_2_ before the measurement. Frozen samples were exposed to excitation wavelength of 436 nm with slit widths of 8 nm for whole cells and 5 nm for isolated complexes in a continuously cooled environment (Morgan-Kiss et al., 2008). For each sample, at-least three replicates of emission spectra were measured.

### PSII fluorescence state transition measurement

Room temperature PSII fluorescence measurements were conducted on cultures induced in state 1 or state 2 as described above. Preliminary analysis was done to identify PSII saturating actinic light intensity and 200 μmol photons m^−2^ s^−1^ was chosen for the subsequent measurements. Log-phase exponentially growing cultures (2 mL) were used. Briefly, measuring light was switched on in the dark and minimal PSII fluorescence (F_O_) was measured. Subsequently, cultures were exposed to 200 μmol photons m^−2^ s^−1^ of actinic red light (λmax=620 nm, 10 Wm^−2^, Scott filter RG 715) to measure maximum fluorescence (F_M_). Percent state transition capacity was calculated using F_M_ values measured under state 1 and state 2 using the formula (F_M_^ST1^ - F_M_^ST2^)/ F_M_^ST1^ % as described before (Girolomoni et al., 2017), where F_M_^ST1^ and F_M_^ST2^ are the maximal PSII fluorescence under state 1 and 2 respectively.

### P700 oxidation-reduction kinetics

Actinic red light induced photooxidation-reduction of P700 was used to determine rates of CEF as previously described (Alric et al., 2010; Morgan-Kiss et al., 2002). Exponential phase cultures (~ 25 μg Chl) were dark adapted for 10 min in presence of DCMU to block electrons from PSII. Dark adapted cultures were then filtered onto 25 mm GF/C filters (Whatman) and measured on the Dual-PAM 100 instrument using the leaf attachment. Absorbance changes at 820 nm were used to calculate proportion of photooxidizable P700, expressed as the parameter ΔA_820_/A_820_. To start the measurement, the signal was balanced and measuring light was switched on. First, P700 was oxidized (P700^+^) by switching on the actinic red light (AL, λmax=620 nm, 10 Wm^−2^, Scott filter RG 715). Subsequently, AL was switched off to re-reduce P700^+^ after steady-state oxidation was reached. The half time for the reduction of P700^+^ to P700 (*t*_½_^red^) was calculated as an estimate of relative rates of PSI-driven CEF (Ivanov et al., 1998). The re-reduction time for P700 was calculated using GraphPad Prism^TM^ software.

### In-vivo spectroscopy measurements

Dark Interval Relaxation Kinetics (DIRK) of electrochromic shift (ECS) were performed on the Kramer Lab IDEA spectrophotometer (Sacksteder & Kramer, 2000; Zhang et al., 2009) to evaluate proton fluxes across thylakoid membrane. Simultaneously, saturation-pulse chlorophyll fluorescence was also measured to estimate the PSII operating efficiency (*ϕ*PSII). Measurements were performed as described before (Kalra et al, 2020). Briefly, 2.5 ml of exponential phase *C. reinhardtii* culture was incubated in presence of bicarbonate under dark condition for 10 min followed by far red exposure for 10 min, to fully oxidize the plastoquinone pool. Cells were incubated in increasing actinic red light for 5 min, and ECS and chlorophyll fluorescence were measured after each light incubation. The PSII operating efficiency was calculated using the formula: (F_M’_-F_S_)/F_M_ and linear electron flow (LEF) was calculated using the equation *A x (fraction_PSII_) x I x ϕPSII* (Baker, 2008), where *A* is absorptivity of the sample (assumed to be 0.84), *fraction_PSII_* is the fraction of light absorbed by PSII stimulating photosynthesis, *I* is the irradiance used and *ϕPSII* is the operating efficiency of PSII as calculated above. In the above equation, *fraction_PSII_* was calculated using the 77K fluorescence spectra and the fraction were 0.56 and 0.579 for low and high salinity conditions, respectively. Proton motif force (*pmf*) was estimated by using the total amplitude of ECS signal (ECS_t_), and the total proton conductivity (*g_H_^+^*) of thylakoid membranes was estimated through the inverse of half time of rapid decay of ECS signal (*τ*_ECS_) during the DIRK measurements (Baker et al, 2007).

### Thylakoid isolation, SDS-PAGE and immunoblotting

Thylakoid membranes were isolated according to previously described protocol (Morgan-Kiss et al., 1998). Briefly, log-phase cells were harvested using centrifugation (2500g at 4°C for 5 min) and resuspended in the cold grinding buffer (0.3 M sorbitol, 10 mM NaCl, 5mM MgCl2, 1mM benzamidine and 1 mM amino-caproid acid). The resuspension was passed through a chilled French press at 10,000 lb/in^2^ two times followed by centrifugation at 23,700g for 30 min to collect thylakoid membranes. The membranes were then washed to remove any impurities using wash buffer (50 mM Tricine-NaOH [pH 7.8], 10 mM NaCl, 5mM MgCl_2_) and the pure thylakoid membranes were collected (13,300 g at 4°C for 20 min). Membranes were resuspended in storage buffer and kept at −80°C until further use.

SDS-PAGE and immunoblotting was performed using 12% Urea-SDS gel (Laemmli, 1970) and as previously described (Kalra et al., 2020). Phosphorylated threonine sites were probed using a primary antibody against P-Thr (Catalog **#** MA5-27976, Thermo Fisher) at 1:500 dilution followed by exposure to protein A conjugated to horseradish peroxidase. Protein blots were detected using ECL Select^TM^ Western Blotting Detection Reagent (Amersham).

### Supercomplex isolation

Sucrose density gradient centrifugation was used to isolate supercomplexes from exponentially grown cultures as previously described (Szyszka-Mroz et al., 2015; Kalra et al, 2020). All steps were performed in darkness and on ice. All buffers contained phosphatase (20 mM NaF) and protease (1 mM Pefabloc SC) inhibitors. Protein complexes were extracted using a 21-gauge needle for further analysis.

### Sample preparation for proteomics

For identifying protein components in the supercomplex, the complex was harvested and 30 μg of total protein was processed for proteomics following the previously published method by Wang et al. (Wang et al., 2016). Samples were digested and cleaned as described before (Kalra et al., 2020).

### Proteomic analyses by liquid chromatography-tandem mass spectrometry (LC-MS/MS)

Two μg of digested peptides were directly loaded onto a capillary C18 column without fractionation and analyzed in a Thermo LTQ Orbitrap XL mass spectrometer. The full mass spectra in the range of 350-1800 m/z were recorded with a resolution of 30,000, and the top 12 peaks of each scan were then selected for further fragmentation for MS/MS analysis. The MS/MS raw data was analyzed using the Patternlab for Proteomic tool (Carvalho et al., 2016). Our UWO241 transcriptomics data was used to generate a UWO241 protein database after supplementing with 37 common contaminants. Reversed sequences were also included as a quality control system to restrain false positive discovery rate to 0.05. *C. reinhardtii* protein database was downloaded from NCBI containing both Swiss-Prot and TrEMBL entries.

## RESULTS

### Salinity tolerance of the three Chlamydomonas species

To identify the salinity sensitivity for all three *Chlamydomonas* species, we conducted a salinity gradient growth experiment. Based on the native habitat of the Antarctic species (ie. the hypersaline Lake Bonney) as well as previous literature on salt tolerance in UWO241 (Pocock et al. 2010) and *C. reinhardtii (*Subramanyam et al. 2010), we selected the following salinity ranges for growth: (i) ICE-MDV and UWO241 were grown in salinity concentrations of 0.43 mM NaCl (BBM medium) to 700 mM NaCl (salinity concentration at 17 m Lake Bonney), and (ii) *C. reinhardtii* was grown in 0.43 mM NaCl to 200 mM NaCl (Subramanyam et al. 2010) (Figure 1). As expected, the halotolerant UWO241 exhibited exponential growth across the full range of salinity conditions. Despite its native environment of hypersalinity, UWO241 exhibited the shortest lag phase and a doubling time of 71.94 ± 2.44 hrs when grown in low salinity (Figure 1 A, D). Growth under the moderate salinity stress of 250 or 500 mM caused a ~1.6-fold increase in doubling time (113 ± 24.13 and 119 ± 3 hrs (p<0.0001), respectively). Last, while the lag phase was significantly longer in UWO241 cultures grown in 700 mM NaCl, once acclimated, these cultures exhibited the fastest doubling time (55 ± 3.38 hr), 1.3-fold faster (p<0.01) relative to BBM-grown cultures (Figure 1, Table 1).

**Figure 1:**
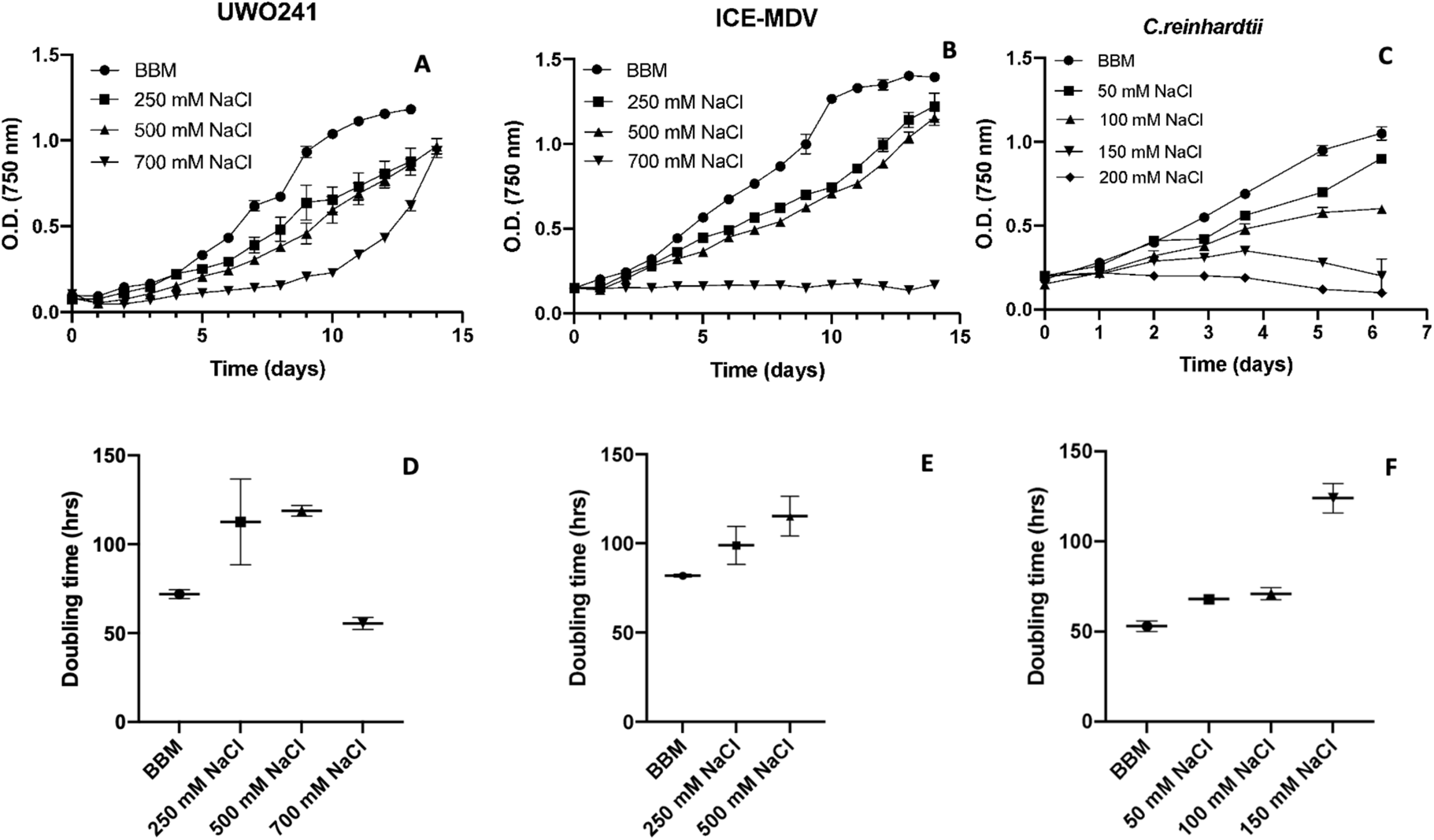
Growth under salinity gradient for the three Chlamydomonas species to identify maximum salinity tolerance. Top Panel - Growth curves, Bottom Panel – Doubling time (A, D UWO241. B, E ICE-MDV. C, F *C. reinhardtii*). Growth was measured as optical density at 750 nm. BBM = 0.46 mM NaCl. (n=3, ±SD).

**TABLE 1.** Physiological characterization of low salt (LS) and high salt (HS) acclimated Chlamydomonas species. Doubling time, photosynthetic parameters and extracted chlorophyll content are shown for all three species (UWO241, ICE-MDV and C. reinhardtii) under the two salinity conditions, low salt (LS) and high salt (HS) (n=3 ± SD). F_v_/F_M_: maximal photosynthetic capacity, YPSII: PSII yield, NPQ: non-photochemical quenching, qP: photochemical efficiency

|  | Doubling time (hr) |  | F<sub>v</sub>/F<sub>M</sub> |  | YPSII |  | qP |  | NPQ |  | Chlorophyll (µg/ml) |  |
| --- | --- | --- | --- | --- | --- | --- | --- | --- | --- | --- | --- | --- |
|  | LS | HS | LS | HS | LS | HS | LS | HS | LS | HS | LS | HS |
| **UWO241** | 71.94<br>±2.44 | 55.46<br>± 3.38 | 0.640<br>±0.008 | 0.585<br>±0.007 | 0.463<br>±0.001 | 0.524<br>±0.029 | 0.719<br>±0.089 | 0.766<br>±0.088 | 0.256<br>±0.026 | 0.307<br>±0.043 | 383.53<br>±18.31 | 221.16<br>±12.69 |
| **ICE-MDV** | 81.88<br>±0.71 | 115.23<br>±11.22 | 0.694<br>±0.009 | 0.669<br>±0.004 | 0.536<br>±0.019 | 0.518<br>±0.007 | 0.733<br>±0.026 | 0.798<br>±0.025 | 0.171<br>±0.017 | 0.223<br>±0.007 | 398.17<br>± 7.43 | 314.50<br>±12.88 |
| **C.**<br>**reinhardtii** | 52.93<br>±2.91 | 70.99<br>± 3.26 | 0.720<br>±0.011 | 0.697<br>±0.019 | 0.621<br>±0.006 | 0.555<br>±0.068 | 0.909<br>±0.060 | 0.829<br>±0.056 | 0.184<br>±0.028 | 0.230<br>±0.037 | 438.26<br>±43.91 | 392.61<br>±54.51 |

ICE-MDV was more recently isolated from Lake Bonney, where it dominates the shallow, freshwater depths (Li et al. 2016; Li and Morgan-Kiss 2019). In the salinity gradient experiment, ICE-MDV grew fastest under control conditions (0.43 mM NaCl, doubling time of 81.88 ± 0.71 hr), but exhibited the ability to grow under a salinity regime of either 250 mM or 500 mM NaCl (doubling times of 98.90 ± 10.68 and 115.23 ± 11.22 hrs, respectively) (Figure 1 B, E). However, unlike UWO241, which has a robust growth at 700 mM NaCl, ICE-MDV was unable to grow in 700 mM NaCl.

The model mesophile *C. reinhardtii* had the highest growth rate under control conditions (doubling time of 52.93 ± 2.91hrs), followed by 50 mM NaCl (doubling time of 67.93 ± 0.38 hrs) (Figure 1 C, F). For *C. reinhardtii,* 100 mM NaCl was the maximum salinity that the cultures exhibited some growth; however, the cultures failed to grow beyond an OD_750_ of 0.6. Last, after a few days of slight growth in the upper salinity levels of 150 mM and 200 mM NaCl, *C. reinhardtii* failed to grow further and entered death phase in both salinity treatments.

Based on these growth physiology results, we chose the following salinity levels for further experiments testing long- and short-term acclimation responses. For low salinity (LS), all strains were grown in BBM medium (0.43 mM NaCl). For high salinity (HS), we used BBM supplemented with, (i) 700 mM NaCl for UWO241, (ii) 500 mM NaCl for ICE-MDV and (iii) 50 mM NaCl for *C. reinhardtii* (Table 1). Cultures were acclimated by serial sub-culturing in the same condition for 14 – 30 days depending upon the growth rate. Photosynthetic and physiological measurements were conducted for all three strains under the two salinity levels (Table 1). All three strains maintained a similar capacity of photosynthesis (F_v_/F_m_, YPSII) after acclimation to the two salinity conditions.

### Cyclic electron flow and PSI activity

We monitored PSI activity in ICE-MDV and *C. reinhardtii*. P700 oxidation/reduction kinetics were conducted on log-phase cultures to monitor changes in P700 photooxidation activity and PSI-mediated CEF in response to long-term salinity acclimation (Figure 2, Figure S1). LS-grown cultures of UWO241 exhibited a *t*_½_^red^ of 259 ± 45 ms. In agreement with previous literature (Kalra et al. 2020; Szyszka-Mroz et al. 2016), acclimation of UWO241 to HS resulted in a 1.6-fold faster re-reduction rate (*t*_½_^red^ = 162 ± 14 ms) (Figure 2 A). Under LS, ICE-MDV exhibited a comparable *t*_½_^red^ value as LS-grown UWO241 (*t*_½_ = 291 ± 42 ms) (Figure 2 B). ICE-MDV responded to HS by a 2.3-fold faster *t*_½_^red^ (*t*_½_^red^ = 124 ± 32 ms). On the other hand, *C. reinhardtii* exhibited a slower *t*_½_^red^ rate in LS media (*t*_½_ = 495 ± 43 ms) which was around 1.6 - 1.9-fold slower than LS-grown ICE-MDV and UWO241 (Figure 2 C). *C. reinhardtii* displayed 1.5 times faster re-reduction rates after acclimation to salinity stress (*t*_½_^red^ = 311 ± 10 ms; Figure 2 C) which matched the salinity response of the psychrophiles. In addition, we also measured change in P_700_ absorbance (ΔA_820_ /A_820_) after AL illumination which reflects the redox state of P700. Both the psychrophiles, (Figure 2 D, E) displayed lower ΔA_820_ /A_820_ values when compared to *C. reinhardtii* (Figure 2 F) (1.5- and 1.7-fold lower, respectively), indicating a reduced capacity for PSI oxidation. Interestingly, the ΔA_820_ /A_820_ values were further reduced significantly in the psychrophiles (Figure 2 D, E) after salinity stress acclimation, while *C. reinhardtii* did not show any significant change in ΔA_820_ /A_820_ after salinity stress acclimation (Figure 2 F).

**Figure 2:**
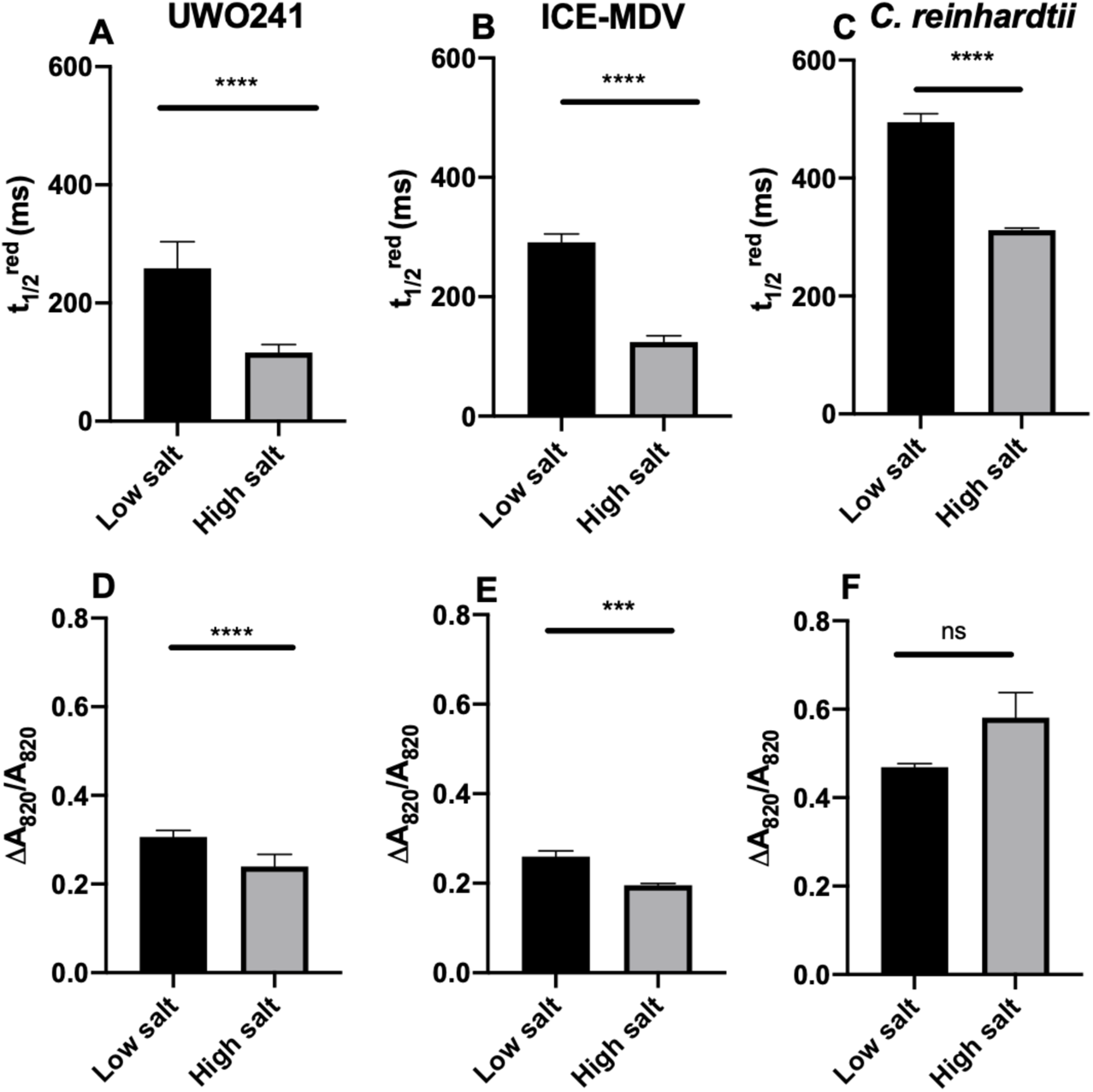
P700 oxidation/reduction analysis on the three *Chlamydomonas* spp. under low and high salinity. Top panel: Re-reduction rate (t_1/2_^red^) was calculated under low (black) and high salinity (grey) for all three strains: UWO241 (A), ICE-MDV (B), *C. reinhardtii* (C). Bottom panel: The proportion of photo-oxidizable P700 is shown as change in absorbance at 820 nm (ΔA_820_/A_820_) for all three strains under low and high salinity: UWO241 (D), ICE-MDV (E), *C. reinhardtii* (F). Actinic red light was used with DCMU to inhibit electron flow from PSII. (*n*=9, ± SD, ns (not significant, p > 0.05), ** (p<0.01), *** (p<0.005), **** (p<0.001))

We also validated our *C. reinhardtii* P700 CEF data with electrochomic shift (ECS) kinetics which estimates light-dependent photosynthesis driven transthylakoid proton flux using IDEA spectrophotometer (Baker et al., 2007) (Figure S3). We used dark interval relaxation kinetics (DIRK) to observe the shift in the electrochomic signal at 520 nm (Kramer et al., 2003). The proton motive force (pmf) was estimated from the total ECS signal (ECS_t_) in *C. reinhardtii* under both low and high salinity conditions (Figure S3 A). We detected higher pmf in HS-acclimated *C. reinhardtii* in all light intensities and the pmf was significantly higher under light intensities of 200 µmol photons m^−2^ s^−1^ and above compared to low salt conditions (~1.6 fold).

This increase in pmf can be attributed to either decrease in ATP synthase activity or proton efflux, or an increase in proton flux through linear electron flow or cyclic electron flow. To identify the processes contributing to increased pmf in HS cultures, we measured the transthylakoid proton conductivity (g_H_^+^) and flux (v_H_^+^) through ATP synthase (Carrillo et al., 2016; Kanazawa & Kramer, 2002; Livingston et al., 2010) (Figure S3 B, C). Proton conductivity or permeability (g_H_^+^) is estimated from inverse of half-time of rapid decay of the ECS signal (*τ*_ECS_) and is dependent on ATP synthase activity (Baker et al., 2007). Both LS and HS cultures showed similar conductivity at lower light intensities, however, at 300 µmol photons m^−2^ s^−1^ and above, the conductivity in HS cultures decreased (~ 1.18 folds) compared to LS cultures (Figure S3 B). On the other hand, the proton flux in HS cultures was consistently higher (1.2 – 2 folds) compared to LS cultures, indicating higher ATP synthesis in HS condition (Figure S3 C). To dissect whether LEF or CEF is contributing to increased proton flux in HS cultures, we plotted v_H_^+^ against LEF. The slope of this curve can inform us about the relative contribution of each pathway towards proton flux (Figure S3 D). We observed that the slope of the HS cultures was 1.25-fold higher than LS cultures, indicating that CEF significantly contributes towards increased proton flux in *C. reinhardtii* HS cultures, which tightly corroborates with our P700 findings. Similar results were also observed for UWO241 cultures acclimated to high salinity in our previous study (Kalra et al., 2020), further emphasizing the importance of CEF in HS acclimation in both non-halophyte and halophyte *Chlamydomonas* species.

### Effect of long-term high-salinity acclimation on short-term state transition response

Previous research showed that UWO241 is a natural state transition mutant (Morgan-Kiss et al. 2002); however, the capacity for state transitions in the sister species ICE-MDV has not been tested. We conducted state transition tests on all three *Chlamydomonas* species after low and high salinity acclimation (Figure 3, Figure S2). In agreement with previous reports, UWO241 cells exhibited very low PSI fluorescence under low salinity, which was further reduced in high salinity (Figure 3 A). State transition treatment had no effect on UWO241 grown under either condition (Figure 3 A).

**Figure 3:**
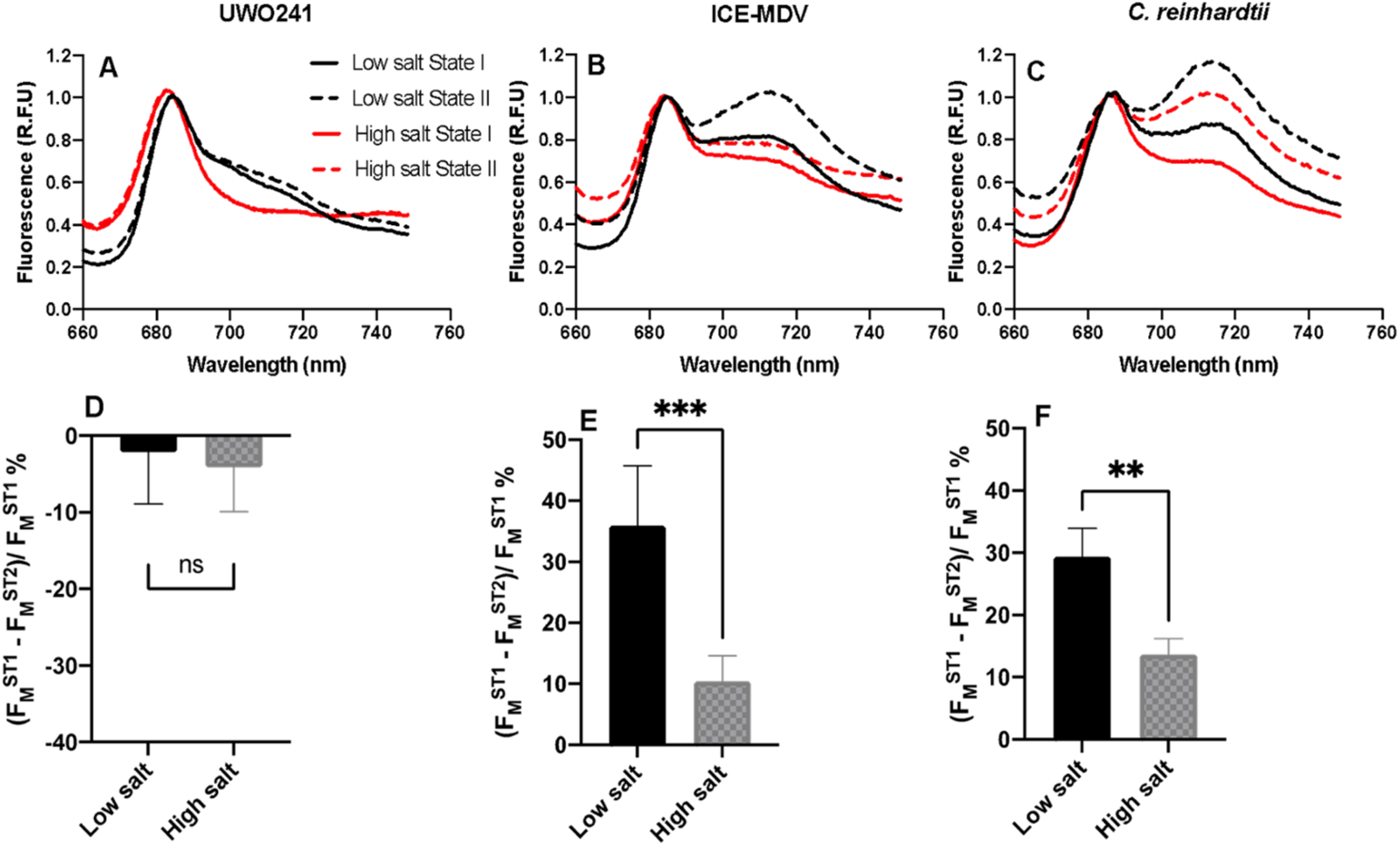
State transition tests after acclimation to low and high salinity in *Chlamydomonas* species. Top panel: Low temperature (77K) fluorescence spectra of the three *Chlamydomonas* spp. under state I and state II conditions after low and high salinity acclimation. Fluorescence values are shown as relative fluorescence units (R.F.U) for each strain: UWO241 (A); ICE-MDV (B); *C. reinhardtii* (C). Low salinity - Black, High salinity - Red. State I – Closed line, State II – dotted line. Bottom panel: Maximal capacity for switching LHCII antenna during State transition induction calculated using room temperature PSII maximum fluorescence (F_M_) as described before (Girolomoni et al., 2017) for each strain: UWO241 (D); ICE-MDV (E); *C. reinhardtii* (F). ST1: state 1, ST2: state 2. (*n*=4-6; ±SD; ns (not significant, p > 0.05), ** (p<0.01), *** (p<0.005))

While ICE-MDV was isolated from the same Antarctic lake as UWO241, its 77K fluorescence emission spectra characteristics were markedly different from that of UWO241 (Figure 3 B). First, under low salinity, ICE-MDV exhibited detectable levels of PSI fluorescence. In addition, when exposed to state 2 conditions, ICE-MDV LS responded with a 1.25-fold increase in PSI fluorescence, suggesting that unlike UWO241, ICE-MDV has retained the ability to undergo state transitions (Figure 3 B). Acclimation to HS resulted in a 1.2-fold reduction PSI fluorescence emission in HS- versus LS-conditions. Furthermore, under state transition conditions, no detectable change in PSI fluorescence was observed in HS-acclimated ICE-MDV cells (Figure 3 B).

As expected, under low salt conditions, *C. reinhardtii* exhibited a typical 77K fluorescence emission spectrum and the ability to undergo a state transition (Figure 3 C). Growth under high salinity resulted in a 1.2-fold decrease in PSI fluorescence yield. Last, unlike the psychrophile strains, *C. reinhardtii-*HS cultures retained the ability to undergo a state transition, exhibiting a comparable response to state 1 conditions as the LS cultures (Figure 3 C).

State transition capacity can also be measured as a loss in maximum fluorescence of PSII (F_M_) at room temperature via detachment of LHCII antenna proteins. We used this measurement to validate the 77K fluorescence emission data as described before (Girolomoni et al., 2017). This test confirmed the 77K fluorescence emission results that UWO241 lacks the capacity for state transitions, regardless of growth conditions (Figure 3 D). In contrast, both ICE-MDV-LS and *C. reinhardtii*-LS cells exhibited 36 and 29 % state transition capacity, which also agreed with the 77K fluorescence emission results. In addition, both ICE-MDV and *C. reinhardtii* exhibited a significant reduction in state transition capacity, following acclimation to high salt (3.5- and 2.2-fold, respectively, relative to LS conditions; p<0.01; Figure 3 E, F).

### Effect of salinity on NPQ capacity and its relationship with CEF

To understand the effect of salinity on NPQ, we measured the NPQ capacity of all the three strains under low and high salinity during a light curve (Figure 4 A). UWO241 displayed increased capacity for NPQ at high salinity at every light level; however, the difference was not significant. On the other hand, *C. reinhardtii* and ICE-MDV showed higher NPQ capacity at high salinity only at the maximum light intensity (830 µmol photons m^−2^). Overall, *C. reinhardtii* displayed highest NPQ capacity, which was significantly higher than UWO241 and ICE-MDV under low salt conditions. Among the psychrophiles, UWO241 had significantly higher NPQ capacity than ICE-MDV under both low and high salinity conditions (Figure xx).

**Figure 4:**
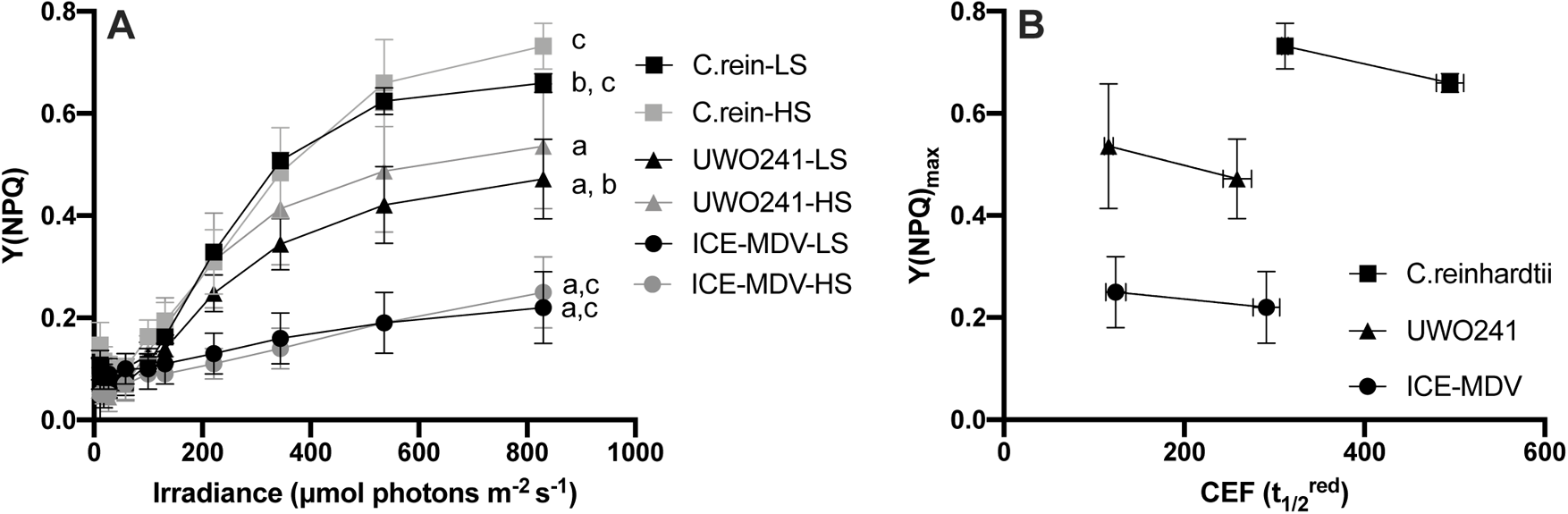
Effect of salinity on non-photochemical quenching (NPQ) capacity and relationship with cyclic electron flow (CEF). (A) NPQ capacity (Y(NPQ)) was measured for the three species during a light curve under low (LS, black) and high salinity (HS, grey) conditions in the three strains *C. reinhardtii* (C. rein, square), UWO241 (triangle) and ICE-MDV (circles). Statistically significant differences between UWO241 vs ICE-MDV (a) and *C. reinhardtii* (b) as well as *C. reinhardtii* vs ICE-MDV (c) are shown (Welch’s t-test, p < 0.05). (B) Relationship between maximum NPQ capacity (Y(NPQ)_max_) and CEF (re-reduction time, t_1/2_^red^) is shown for the two salinity conditions for all species.

Next, we evaluated the relationship between maximum NPQ capacity and CEF for the three species (Figure 4 B). As expected, increase in NPQ was corelated with higher CEF (i.e. faster t_1/2_^red^) in all three species but the slope of this increase varied with organism. The slope of increase was highest in UWO241 (4.5×10^−3^ units?) followed by *C. reinhardtii* (3.9 x10^−3^) and last ICE-MDV (1.8 x10^−3^).

### Thylakoid protein phosphorylation

In *C. reinhardtii,* several key photosynthetic proteins are phosphorylated in response to changes in environmental conditions. State transitions are accompanied by transient phosphorylation of LHCII proteins through STT7 kinase (Lemeille & Rochaix, 2010). Previous reports have shown that the thylakoid proteome of UWO241 is under-phosphorylated relative to other photosynthetic organisms and exhibits novel high molecular weight phospho-proteins (Morgan-Kiss et al. 2002; Szyszka et al. 2007). We compared thylakoid phospho-protein profiles of the three strains acclimated to either LS or HS (Figure 5). When probed with a phospho-threonine antibody, phosphorylation of major LHCII proteins was not observed in UWO241 grown under either LS or HS conditions (Figure 5, A). The phosphoprotein profile of thylakoids from UWO241 exhibited phosphorylation of several high molecular weight proteins (~150 and 250 kDa) under both LS and HS.

**Figure 5:**
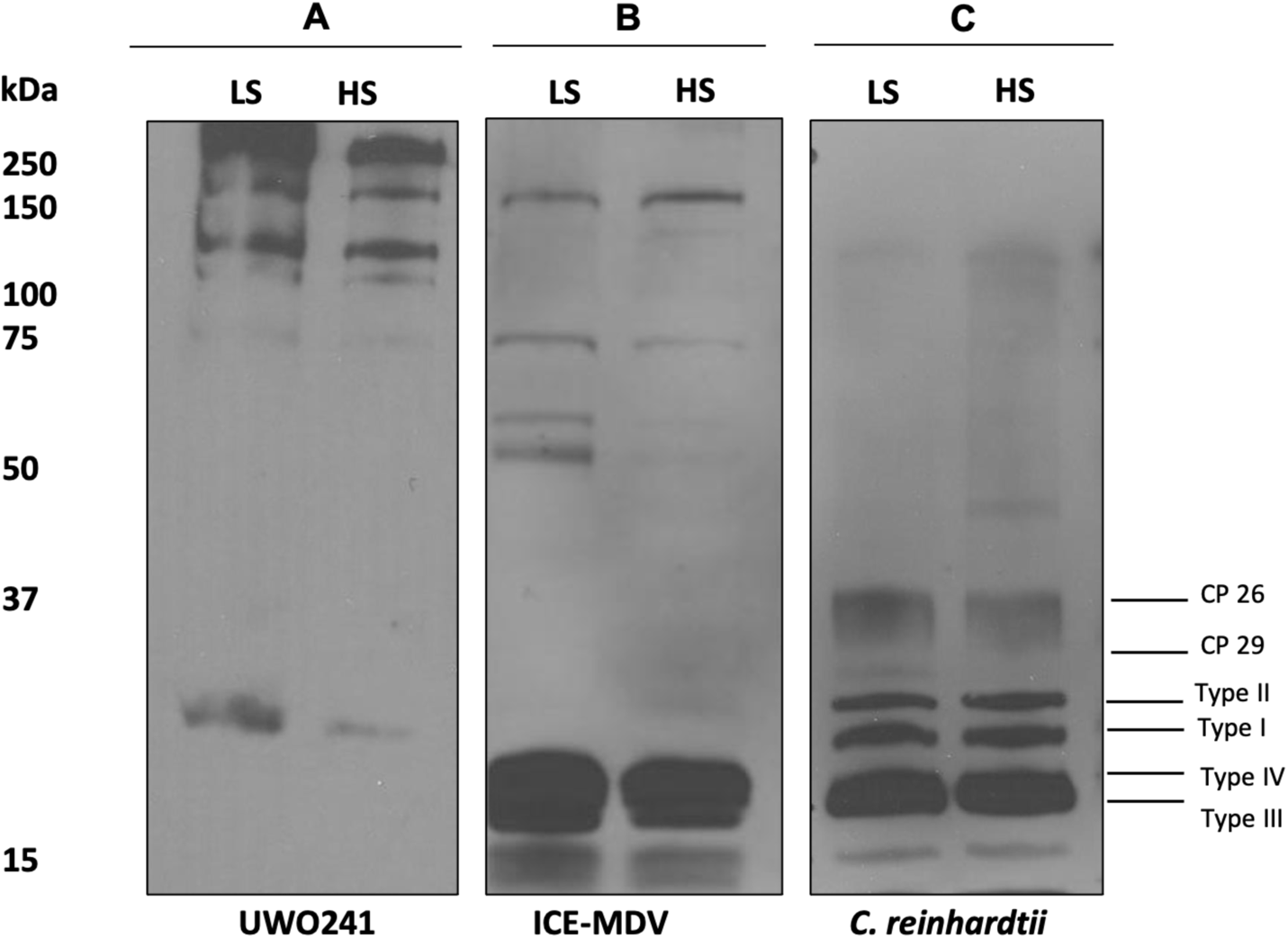
Thylakoid phosphorylation pattern of the three *Chlamydomonas* spp. under low (LS) and high (HS) salinity. Isolated thylakoids from three species were run on separate 12% SDS-PAGE and probed with phospho-threonine antibody. Panel A: UWO241, Panel B: ICE-MDV, Panel C: *C. reinhardtii*. Mol. wt ladder (KDa) is shown on the left. The different LHCII types are labelled on the right.

ICE-MDV exhibited phosphorylation of some major LHCII proteins (type III and IV) under either LS or HS; however, some phospho-LHCII proteins that were detected in *C. reinhardtii* were not detectable (Figure 5, B). Last, thylakoids isolated from LS- or HS-grown ICE-MDV also possessed several high molecular weight phospho-proteins (~150 kDa), albeit at lower levels compared with UWO241. *C. reinhardtii* exhibited phosphorylation of several thylakoid proteins, specifically phosphorylated LHCII (type I, II, III and IV) (Figure 5, C). Growth in high salinity did not alter the pattern or abundance of phosphoproteins of *C. reinhardtii.* Last, we did not detect phosphorylation of larger proteins in *C. reinhardtii* under either growth condition.

### Assembly of chlorophyll protein supercomplexes under high salinity

PSI-supercomplexes have been shown to be associated with conditions promoting CEF. In UWO241, formation of a stable PSI-supercomplex is proposed to be essential for maintaining sustained high rates of CEF (Szyska-Mroz et al. 2015; Kalra et al. 2020). We conducted sucrose density gradient centrifugation on solubilized thylakoid membranes from the three *Chlamydomonas* species after acclimation to salinity stress. As a control, we separated Chl-protein complexes from solubilized thylakoids from *C. reinhardtii* exposed to either state 1 or state 2. Under State 1, solubilized thylakoids of LS-grown *C. reinhardtii* exhibited three Chl-protein bands: (i) trimeric LHCII (band 1), (ii) PSII core (band 2) and (iii) PSI-LHCI complex (band 3) (Figure 6 A). As expected, State 2 treated-thylakoids exhibited a fourth band representing the PSI-supercomplex (Figure 6 A, band 4).

**Figure 6:**
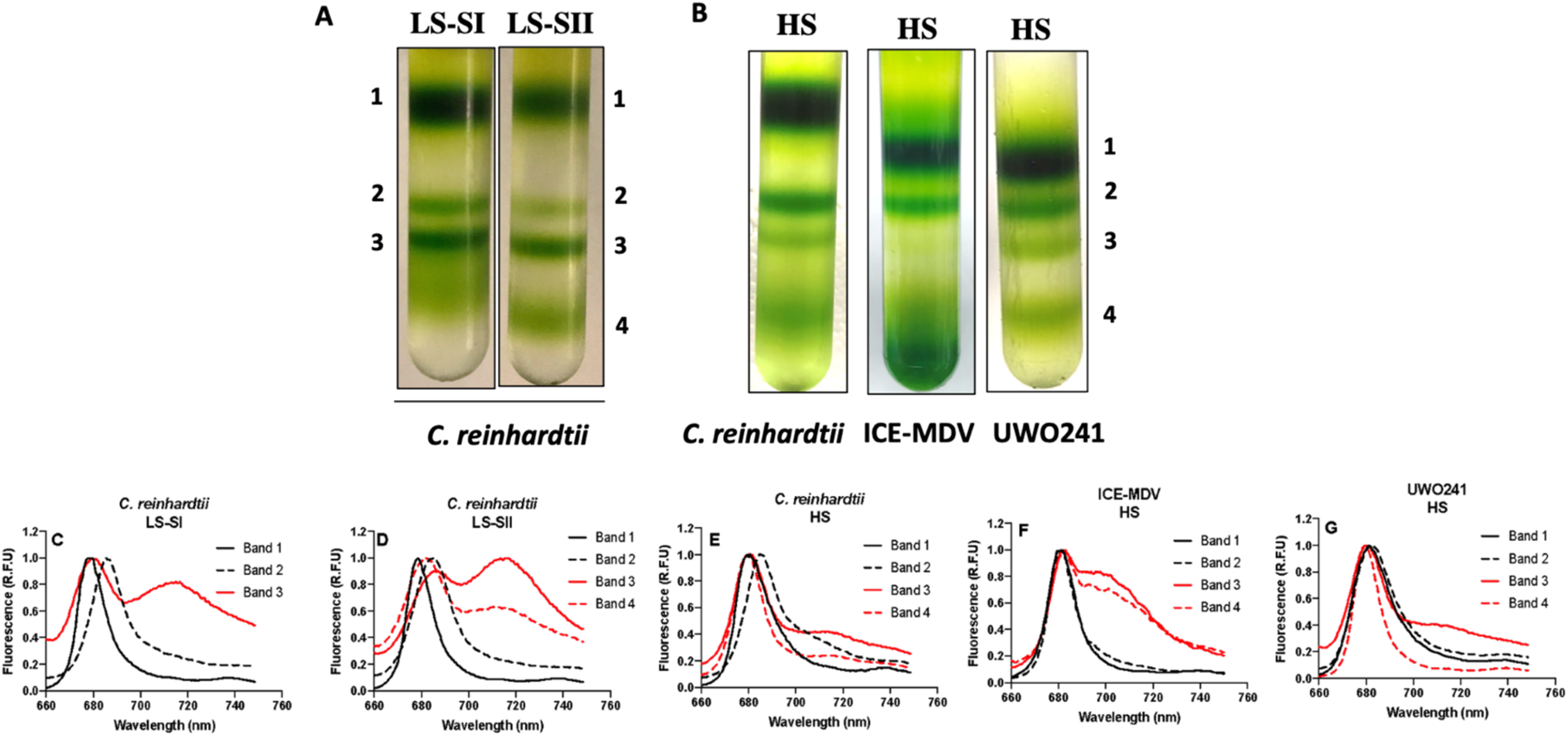
Isolation of supercomplexes from conditions promoting CEF in *Chlamydomonas* species. Top panel: Separation of protein complexes on a sucrose density gradient for A. model mesophile *C. reinhardtii* during state transitions and B. model mesophile and the psychrophiles ICE-MDV and UWO241 under high salinity. Bottom panel: 77K fluorescence spectra for protein complex bands isolated from sucrose density gradient: *C. reinhardtii* under low salt and state 1 (C), state 2 (D), under high salinity (E); ICE-MDV under high salinity (F); UWO241 under high salinity (G). LS-SI: Low salinity, state 1; LS-SII: Low salinity, state 2; HS: High salinity. Band 1: LHCII, Band 2: PSII, Band 3: PSI-LHCI, Band 4: Supercomplex.

Next, we compared the banding patterns in sucrose density gradients from thylakoids isolated from all three *Chlamydomonas* species after acclimation to salinity. In response to HS, all strains exhibited a reduction in the relative levels of PSI (Band 3). In addition, thylakoids isolated from HS-acclimated cultures of all three *Chlamydomonas* species exhibited the appearance of the supercomplex band (band 4; Figure 6 B).

Bands from the gradients were collected for 77K Chl fluorescence analysis (Figure 6C-G). In *C. reinhardtii* exposed to short-term conditions, emission peaks for the major Chl-protein complexes agreed with previous reports (Iwai et al., 2010). In state 1, Band 1 and 2 exhibited major fluorescence peaks at 678 nm (LHCII) and 682 nm (PSII core) respectively, while Band 3 exhibited emission peaks at both 678 nm and 712 nm (PSI-LHCI) (Figure 6 C). In state 2, Band 4 exhibited a similar emission spectrum pattern as Band 3, with a reduction in 712 nm emission (Figure 6 D).

The 77K fluorescence emission spectra patterns of the PSI and supercomplex bands exhibited strain- and growth condition-distinctions (Figure 6 A, B). First, HS acclimation in *C. reinhardtii* resulted in a loss of PSI fluorescence emission at 712 nm in Bands 3 and 4 (Figure 6 E). In contrast, ICE-MDV-HS thylakoids exhibited a broad shoulder in fluorescence emission between 705 - 712 nm for Bands 3 and 4. In agreement with Kalra et al. (2020), HS-acclimated UWO241 exhibited minimal PSI fluorescence for Bands 3 and 4 (Figure 6 G).

### Proteome analysis of high salinity-associated supercomplexes

Recent reports have shown that the HS-associated supercomplex of UWO241 exhibits some distinct compositional features compared with supercomplexes isolated from other algae, including the presence of novel proteins and a depletion in LHCI and LHCII polypeptides (Kalra et al. 2020). We compared SC composition in *C. reinhardtii* and UWO241 after acclimation to high salinity (HS) with *C. reinhardtii* under state 2 conditions (SII), using shotgun proteomics (Figure 7; Table 2). All three supercomplexes had proteins of PSI, LHCI, LHCII and Cyt b_6_f (Table 2). However, relatively higher number of polypeptides belonging to these three thylakoid complexes were identified in the supercomplexes from *C. reinhardtii* compared to the other two species. For example, the *C. reinhardtii* supercomplexes contained PSI core subunits while UWO241-HS supercomplex was missing several, including PsaA and PsaB. However, using western blotting, we could detect PsaA in the supercomplex band (Band 4, Figure S4) as well as PSI-LHCI complex (Band 3, Figure S4). Both *C. reinhardtii* supercomplexes contained all subunits of Cyt b_6_f, whereas only core subunits PetA, PetB, and PetC were detected in UWO241.

**Figure 7:**
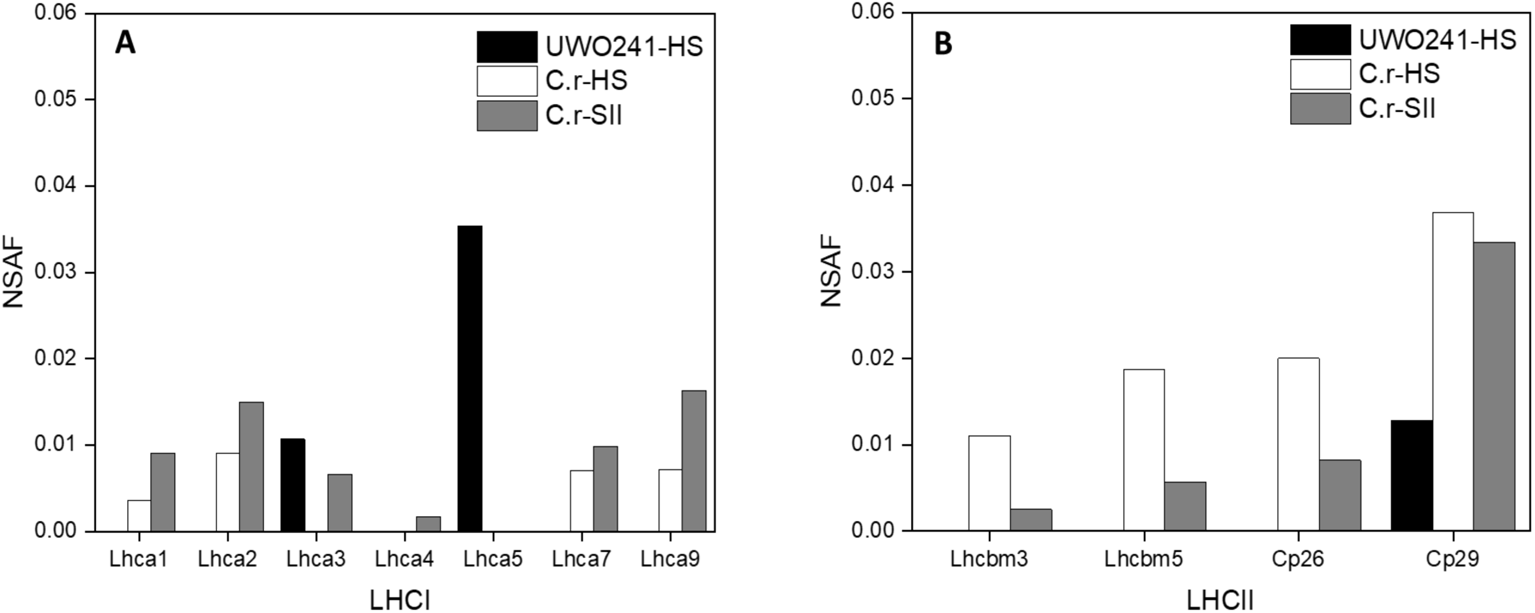
Proteome comparison of light harvesting complexes of supercomplex fractions from *C. reinhardtii* and UWO241. Protein composition of *C. reinhardtii* supercomplexes isolated under state 2 and after high salinity acclimation, as well as UWO241 supercomplex after high salinity acclimation are shown. The normalized spectral abundance factor (NSAF) for each identified protein within a supercomplex were calculated to compare the relative abundance of subunits across species and treatments. The major light harvesting complexes and their subunits participating in supercomplex formation are shown here: Light Harvesting complex I (LHCI, A), Light Harvesting complex II (LHCII, B). UWO241-HS (UWO241-HS, Black), *C. reinhardti*i-HS (C.r-HS, White), *C.reinhardtii* state 2 (C.r-SII, Grey).

**TABLE 2.**
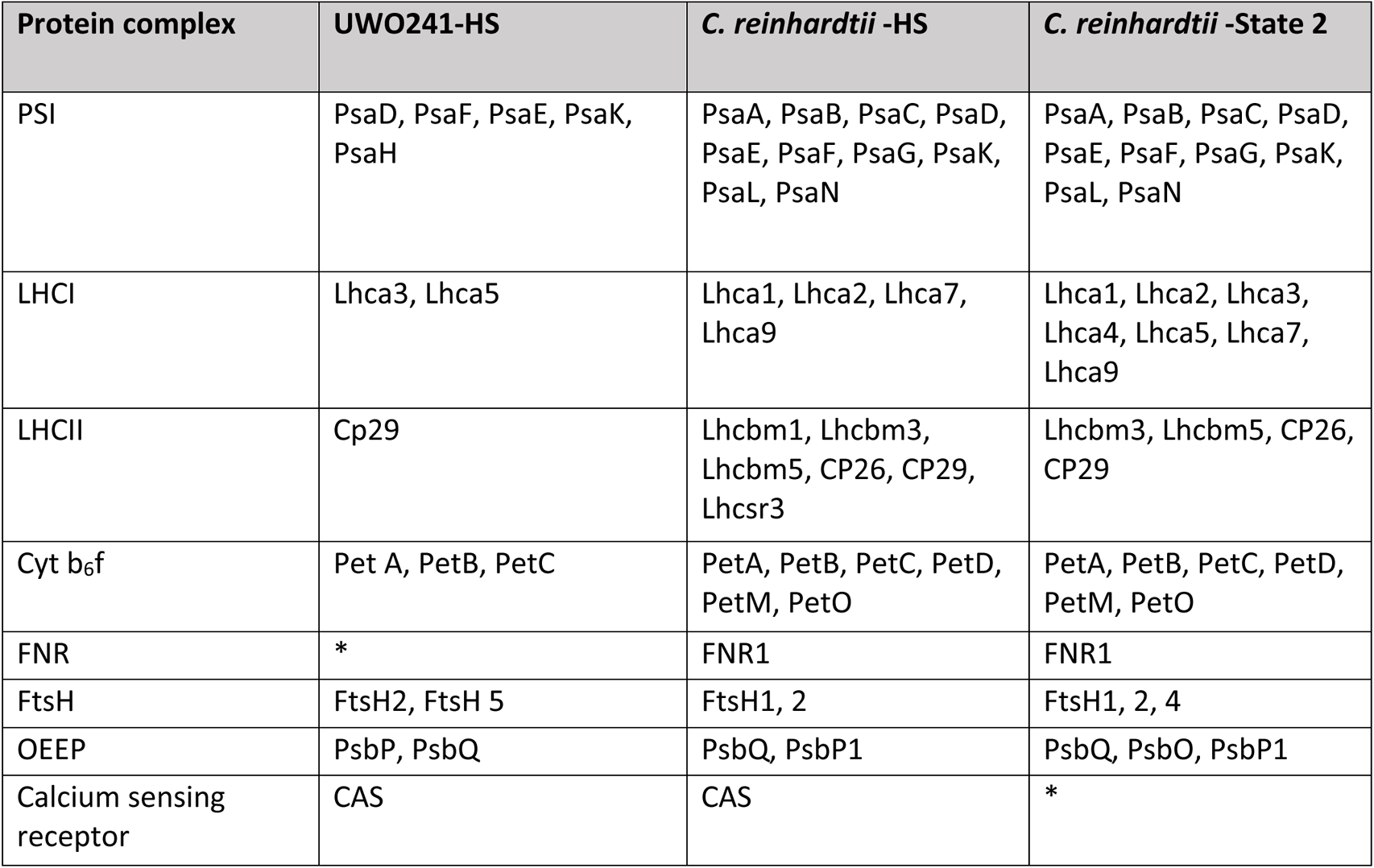
Major proteins involved in supercomplex formation in C. reinhardtii and UWO241. The subunits of each protein identified through shot-gun proteomics are shown. HS: High salinity acclimated; State 2: State 2 locked culture; PSI: photosystem I; LHCI: Light harvesting complex I; LHCII: Light harvesting complex II, Cyt b_6_f: cytochrome b6f, FNR: Ferredoxin NADP reductase, FtsH: ATP dependent zinc metalloprotease, OEEP: Oxygen evolving enhancer protein, CAS: Calcium sensing protein

| Protein complex | UWO241-HS | *C. reinhardtii* -HS | *C. reinhardtii* -State 2 |
| --- | --- | --- | --- |
| PSI | PsaD, PsaF, PsaE, PsaK, PsaH | PsaA, PsaB, PsaC, PsaD, PsaE, PsaF, PsaG, PsaK, PsaL, PsaN | PsaA, PsaB, PsaC, PsaD, PsaE, PsaF, PsaG, PsaK, PsaL, PsaN |
| LHCI | Lhca3, Lhca5 | Lhca1, Lhca2, Lhca7, Lhca9 | Lhca1, Lhca2, Lhca3, Lhca4, Lhca5, Lhca7, Lhca9 |
| LHCII | Cp29 | Lhcbm1, Lhcbm3, Lhcbm5, CP26, CP29, Lhcsr3 | Lhcbm3, Lhcbm5, CP26, CP29 |
| Cyt b<sub>6</sub>f | Pet A, PetB, PetC | PetA, PetB, PetC, PetD, PetM, PetO | PetA, PetB, PetC, PetD, PetM, PetO |
| FNR | * | FNR1 | FNR1 |
| FtsH | FtsH2, FtsH 5 | FtsH1, 2 | FtsH1, 2, 4 |
| OEEP | PsbP, PsbQ | PsbQ, PsbP1 | PsbQ, PsbO, PsbP1 |
| Calcium sensing receptor | CAS | CAS | * |

Notably, the UWO241 supercomplex was depleted for most LHCII and LHCI proteins compared with both supercomplexes isolated from *C. reinhardtii*. The relative abundance of several proteins belonging to light harvesting complexes I and II, which have been previously associated with other supercomplexes, was compared using normalized spectral abundance factor (NSAF) (Figure 7 A, B). LHCI subunit abundance was variable among the different supercomplexes. While *C. reinhardtii*-SII supercomplex contained all but one Lhca subunit, supercomplexes from HS-*C. reinhardtii* and UWO241only contained 4 and 2 Lhcas, respectively (Figure 7 B). Both supercomplexes from *C. reinhardtii* contained the 4 major LHCII subunits. In contrast, the UWO241 supercomplex lacked all LHCIIs except cp29.

## DISCUSSION

Much of the research on acclimation of photosynthetic organisms to environmental stress has intensively focused on a handful of model organisms. There is a significant knowledge gap in identifying acclimation strategies in photosynthetic organisms adapted to permanent stress in their native habitats. In this study, we exploited the salinity tolerance variability of three *Chlamydomonas* spp. to delineate the roles of CEF and PSI-supercomplex formation during long-term stress. *C. priscuii* UWO241 is a well-studied psychrophilic halophyte and exhibited robust growth and photosynthetic activity at 700 mM NaCl. Whilst from the same Antarctic lake, the sister species *C.* sp. ICE-MDV showed lower halotolerance, growing slowly in salinity levels at or above 500 mM NaCl. We attribute the differential salinity sensitivities between the two Lake Bonney algae to the permanent halocline in the water column (Priscu & Spigel, 1996). ICE-MDV dominates the freshwater surface layers, while UWO241 was isolated from the deep hypersaline layers (Neale and Priscu 1995). Thus, these extremophilic green algal strains provide an opportunity to compare the consequences of adaptation to differential long-term salinity stress. We included the salt-sensitive model green alga, *Chlamydomonas reinhardtii*, to delineate between common and novel mechanisms of long-term stress acclimation in salt-sensitive vs. - tolerant *Chlamydomonas* spp.

While CEF is essential in plants and algae for balancing ATP/NADPH (Kramer & Evans, 2011; Lucker & Kramer, 2013) and photoprotection in both PSII (Joliot & Johnson, 2011; Kukuczka et al., 2014) and PSI (Huang et al., 2013; Huang et al., 2017; Yamori & Shikanai, 2016), it has been generally associated with response to short-term, transient stress (Iwai et al. 2010; Takahashi et al. 2013; Strand et al. 2015). In contrast, UWO241 maintains sustained high rates of CEF (Szyszka-Mroz et al. 2015; Kalra et al. 2020). We wondered is this phenomenon an oddity of one extremophilic strain or could high CEF represent a generalized long-term stress strategy? First, we showed that under nonstress conditions both the Antarctic lake algae, UWO241 and ICE-MDV, exhibit markedly faster CEF rates relative to *C. reinhardtii*. More importantly, all three strains responded to long-term salinity stress by increasing CEF; although, CEF rates in *C. reinhardtii*-HS were still significantly slower relative to the extremophiles. Sustained CEF has also been recently reported in other extremophilic algae living in high latitude environments, such as the snow alga *Chlamydomonas nivalis* (Young & Schmidt, 2020; Zheng, Xue et al., 2020). ECS measurements in *C. reinhardtii* and UWO241 higher CEF in high salt-acclimated cells, which was associated with increased proton flux through ATP synthase in both species (Figure S3; Kalra et al., 2020). Increased CEF in all three species was also associated with higher NPQ capacity (Figure 4). We propose that increased CEF can contribute to excess ATP production and quenching of excess energy under long-term salinity stress.

Salinity acclimation involves strain-specific changes in PSI structure and function. One of the early discoveries in UWO241 was that it exhibits permanent downregulation of PSI across a broad range of treatments and growth conditions (Cook et al., 2019; Morgan-Kiss et al., 2002a,b; Morgan et al., 1998). Morgan et al. (1998) and Kalra et al. (2020) linked the constitutively reduced PSI low temperature fluorescence emission in UWO241 with loss of most LHCI polypeptides. A recent report also showed that ICE-MDV and *C. reinhardtii* modulated PSI fluorescence emission in response to iron availability, while UWO241 exhibited minimal changes in PSI (Cook et al. 2019). In agreement with Cook et al. (2019), ICE-MDV exhibited PSI functional characteristics that more closely match *C. reinhardtii* (Figure. 3 A-C). Thus, under low salinity conditions, PSI structure appears to be distinct between the Lake Bonney algae, UWO241 and ICE-MDV. This would be an advantage for ICE-MDV which resides in the more variable, less extreme habitat of the shallow layers of Lake Bonney. During high salt acclimation, all three strains exhibited reduced PSI fluorescence in either whole cells or isolated PSI complexes (Figures. 3 A-C and 5 E-G). These results suggest a common long-term stress-induced effect on PSI organization.

Sustained CEF is associated with formation of a PSI-supercomplex. A high molecular weight band that migrated lower than PSI-LHCI bands was detected in the sucrose density gradients from high salt-acclimated cultures of all three of the *Chlamydomonas* species (Figure 6 B). A PSI-supercomplex was first reported in *C. reinhardtii* when the presence of several LHCII subunits that migrate from PSII to PSI during state 2 transition (Takahashi et al., 2006). Later studies identified additional protein participants in the supercomplex. The state 2 supercomplex was shown to be calcium-regulated, containing CAS, ANR1 and PGRL1 proteins (Terashima et al. 2012). Further research discovered that cyt b_6_f is also essential for supercomplex formation (Minagawa, 2016). A recent structural study identified two LHCI (Lhca2 and 9) subunits whose dissociation is important for PSI-LHCI-cyt b_6_f supercomplex formation (Steinbeck et al., 2018). In addition, a PSI-LHCI-LHCII supercomplex of *C. reinhardtii* under state 2 has two LHCII trimers and ten LHCI subunits (Z. Huang et al., 2021). Our first clue that the HS-induced supercomplexes could be functionally distinct from that of the *C. reinhardtii* State 2 supercomplex came from 77K fluorescence emission spectra of the isolated pigment-protein complexes. Both PSI and the supercomplex isolated from *C. reinhardtii* cells in State 2 exhibited fluorescence emission bands at 720 nm, indicative of PSI fluorescence. In contrast, PSI and supercomplex bands collected from C. *reinhardtii*-HS cells exhibited minimal PSI fluorescence and resembled the emission spectra of UWO241 (Figure 6 D, E, G), indicating the absence of LHCI in the supercomplex.

PSI-supercomplexes are ubiquitous; however, protein composition is strain-specific and dependent upon the time scale of stress exposure. Under transient conditions, several proteins were identified in the *C. reinhardtii* PSI-supercomplexes, including PSI core proteins, cyt b_6_f, LHCI, LHCII, CAS (Calcium Sensing protein), FNR1 (Ferredoxin NADP Reductase), ANR1 (Anaerobic response 1 protein) (Iwai et al., 2010; Steinbeck et al., 2018; Takahashi et al., 2006; Terashima et al., 2012). However, composition of supercomplexes operating under longer term time scales is not known. To further understand the strain- and treatment-specific differences between the supercomplexes, we analyzed the supercomplex components using mass spectrometry. In agreement with previous studies (Z. Huang et al., 2021; Iwai et al., 2010; Steinbeck et al., 2018; Terashima et al., 2012), both the HS and state 2 (SII) *C. reinhardtii* supercomplexes contained 10 out of 13 PSI subunits. In stark contrast, the UWO241 supercomplex only contained 4 core PSI proteins. The supercomplexes also contained varying numbers of Lhca proteins. Relative to the SII supercomplex, the HS supercomplex of *C. reinhardtii* was missing three Lhca proteins (Lhca3, Lhca4, Lhca5). In agreement with Kalra et al. (2020), the UWO241-HS supercomplex lacked all Lhca proteins except Lhca3 and Lhca5. Thus, reduction in Lhca proteins and a reduced PSI peak in the supercomplex 77K fluorescence emission spectra appear to be part of salinity acclimation in *C. reinhardtii*. Recently, cyt b_6_f has been shown to be an important member of the state 2 supercomplex, where electron transfer activity revealed reduction of cyt b subunit through ferredoxin (Minagawa, 2016). In our study both HS supercomplexes of UWO241 and *C. reinhardtii* contained several cyt b_6_f subunits; however, UWO241 supercomplex had disproportionately higher abundance of core Pet A, B and C subunits, and only trace levels of the other Pet proteins (Table 2).

Presence of a salt-associated supercomplex was associated with reduced state transition capacity. State transitions are short-term acclimatory mechanisms that re-balance excitation energy under conditions of an over-reduced PETC and are shared across broadly diverse photosynthetic lifeforms (Wollman, 2001). UWO241 is a state transition mutant which does not phosphorylate major LHCII (Morgan-Kiss et al., 2002). Even though ICE-MDV has evolved in the same lake and is also a psychrophile, ICE-MDV exhibited a typical capacity for state transitions which was comparable with that of *C. reinhardtii* (Figure 3 B). Thus, a loss of state transitions is not a consequence of psychrophily. Instead, when ICE-MDV and *C. reinhardtii* were acclimated to high salinity, both strains exhibited significant losses in state transition capacity (Figure 3 E, F). These results suggest that long-term acclimation to salinity and formation of a supercomplex attenuates the capacity for state transitions. This phenomenon is further enhanced during adaptation to a hypersaline environment to the extent that the mechanism was permanently lost in UWO241. Takizawa et al. (2009) also observed that LS-grown cells of UWO241 were sensitive to oxidizing or reducing conditions of DCMU (state 1) and FCCP (state 2), respectively, while HS-grown cultures were remained locked in State 1. More recently, Szyzska-Mroz et al. (2019) reported that UWO241 may utilize a poorly understood spill-over mechanism instead of classic state transition; although, the localization of PSII and PSI in UWO241 thylakoids is unknown and an older report suggested that the two complexes are not close (Morgan-Kiss et al. 2002).

Adaptation to low temperatures and high salinity has led to differential thylakoid phospho-protein patterns. The aberrant capacity for state transitions in UWO241 was previously linked with to an inability to phosphorylate LHCII polypeptides based on immunoblotting with phospho-threonine antibodies (Morgan-Kiss et al. 2002a). Instead, novel, high molecular mass phosphoproteins of >130 kD as well as a 17 kD polypeptide identified as a PsbP-like protein were identified in the thylakoids and supercomplexes isolated from HS-grown UWO241 cells (Szyszka-Mroz et al. 2015). More recently, Szyzska-Mroz et al. (2019) reported that UWO241 does exhibit light-dependent [γ-33P] ATP labeling of thylakoid polypeptides, including limited phosphorylation of LHCII proteins. The phospho-protein patterns were unique in UWO241 compared with *C. reinhardtii*, and phosphorylation required low temperatures (Szyzska-Mroz et al. 2019). In the current study, we confirmed the unique phosphorylation patterns of UWO241 thylakoids relative to *C. reinhardtii*, with minimal phosphorylation of LHCII and the appearance of multiple high molecular weight phospho-proteins. In contrast, phospho-protein pattens in ICE-MDV exhibit features of both UWO241 and *C. reinhardtii*, with the presence of major LHCII phospho-proteins and the appearance of higher molecular weight bands (Figure 5). These differences between UWO241 and ICE-MDV fit well with the retainment of state transition ability in ICE-MDV.

What structural or functional alterations in the PETC associated with long-term salinity acclimation could be contributing to altered state transition response? A major consequence of a state transitions is formation of PSI-LHCI-LHCII supercomplex and higher CEF (Iwai et al., 2010). Takahashi et al. (2013) elucidated that although both CEF and state transitions are controlled through redox status of the plastoquinone pool, they can occur independent of each other. In UWO241 a restructured photosynthetic apparatus that is primed for constitutive capacity for CEF is key in high salinity acclimation (Szyszka-Mroz et al. 2105; Kalra et al., 2020). We surmised that assembly of stable PSI-supercomplexes could be a general acclimatory response to deal with long-term high salinity. Restructuring of the photosynthetic apparatus to provide sustained CEF, may inhibit the state transition response.

In contrast with our findings that acclimation and adaptation to salinity stress interferes with state transition capacity in *Chlamydomonas* species, the model halophile *Dunaliella* is capable of state transitions (Li et al., 2019; Petrou et al., 2008). State transition ability in *Dunaliella* appears to be associated with different configurations of PSI. Perez-Boerema et al. isolated a minimal PSI from *D. salina* which is missing several PSI core subunits which are necessary for state transitions (Perez-Boerema et al., 2020). The structure of the ‘Basic PSI’ had only 7 core subunits (PsaA-F; PsaJ). Caspy and colleagues then isolated a ‘Large PSI’ containing additional core subunits, including PsaL and PsaO (Caspy et al., 2020). The authors suggest that small and large PSI conformations allow green algae to modulate the function of PSI in variable environments. Our findings extend these recent structural studies by linking PSI-supercomplexes with PSI function. The salt-tolerant UWO241 supercomplex appears to possess the small conformation of PSI, while the salt-sensitive *C. reinhardtii* supercomplex appears to contain the larger PSI.

While it is often assumed that global climate change is mainly associated with increasing temperatures, this is a simplistic view. In fact, the direct and indirect effects of environmental change on the growth and productivity of photosynthetic organisms residing in different habitats is complex. As the climate change exacerbates, there is a growing need for an improved understanding of how organisms will respond to and survive a myriad of stress conditions, especially long-term steady-state stresses (Alexandratos & Brunismas, 2012). The function of the photosynthetic apparatus is key to survival under environmental change: CEF is an essential pathway in almost all photosynthetic organisms, making it an ideal candidate to study stress acclimation in the context of climate change (Kramer & Evans, 2011). Understanding how CEF can help plant and algal survival under physiologically relevant, steady-state stress conditions can help us engineer photosynthetic organisms to better withstand climate change in the future.

## CONCLUSIONS

We show here that sustained CEF supported by restructuring of PSI and formation of a supercomplex is an important strategy in green algae to deal with long-term high salinity stress. CEF has the dual benefit of providing photoprotection of both PSI and PSII and balancing energy needs. Our study suggests that green algae adapted to different environmental stressors have evolved to activate CEF and titer the stability of the PSI-supercomplex to support stress responses over broad time scales. Under short-term stress, state transitions and reversible phosphorylation of LHCIIs mediate formation of the transient PSI-LHCI-LHCII supercomplex. During longer time scales when organisms need to fully acclimate, formation of a stable PSI-supercomplex to support sustained levels of high CEF is essential. Under the long-term stress conditions, state transition capacity is transiently lost until the stress event dissipates. Last, in extremophiles which exhibit adaptation under permanent abiotic stress, evolution of constitutive CEF and additional changes to the supercomplex to further enhance stability helps maintain robust growth and photosynthesis, but at the expense of full loss of state transitions. Further research is needed to better understand the stability and functional differences between the green algal PSI conformations.

## SUPPLEMENTARY DATA

The following supplementary data are available:

**Supplementary Figure S1: P700 reduction kinetics of the three *Chlamydomonas* species under low and high salinity**. A. UWO241, B. ICE-MDV, C. *C. reinhardtii.* Low salinity (LS) traces are shown in blue. High salinity (HS) traces are shown in red.

**Supplementary Figure S2: PSII state transition test for the three *Chlamydomonas* spp under low (LS) and high (HS) salinity.** The maximum PSII fluorescence values (F_MAX_) are shown for all three strains (UWO241-A, ICE-MDV-B and *C. reinhardtii*-C) under state I and state II conditions. State I: DCMU, State II: FCCP. (*n*=4-6, dotted line=mean value)

**Supplementary Figure S3: Electrochomic shift (ECS) measurements for *C. reinhardtii* after low and high salinity acclimation.** A. Total proton motive force (pmf) is shown as total change in the ECS signal (ECS_t_) for low (LS) and high (HS) salt acclimated cultures under increasing light intensities. B. ATP synthase conductivity (g_H_^+^) C. Total proton flux (v_H_^+^) and D. Change in total proton flux as a function of linear electron flow (LEF). (*n*=3, ± SD, * (p < 0.05), ** (p<0.01), *** (p<0.005), **** (p<0.001))

**Supplementary Figure S4: Immunoblot of PsaA in UWO241-HS.** Whole thylakoids and protein complex fractions collected from sucrose density gradient centrifugation (Bands 1, 3 and 4) were run on SDS PAGE and probed using psaA antibody. Molecular wt ladder is shown on left.

## Abbreviations

CEF: cyclic electron flow
PETC: photosynthetic electron transport chain
PSI: photosystem I
PSII: photosystem II
LHCI: light harvesting complex I
LHCII: light harvesting complex 2
Cyt b_6_f: cytochrome b_6_f
NPQ: non-photochemical quenching
FNR: ferredoxin NADP reductase
ANR1: anaerobic response I protein
CAS: calcium sensing protein
PGRL1: proton gradient like protein 1
ECS: electrochomic shift
DIRK: dark interval relaxation kinetics

## ACKNOWLEDGEMENTS

We thank members of Kramer laboratory for their valuable suggestions and inputs on IDEA spectrophotometer optimization. We also acknowledge Will McHargue for his help with IDEA spec measurements and bioinformatic processing of fluorescence data. Last, we would like to thank Microbiology Department at Miami University for the instrument support for Mass spectrometry.

## AUTHOR CONTRIBUTIONS

I.K. and R.M.K. conceptualized the study design. I.K. conducted the photobiology, physiology, spectroscopic and proteomics experiments. R.M.K. supervised the entire study. R.Z provided IDEAspec support and supervised the measurements. X.W. ran and analyzed the proteomics data. I.K and R.M.K wrote the original article. I.K, R.M.K, R.Z and X.W edited the final article.

