## Supplementary_figures for "High salinity activates CEF and attenuates state transitions in both psychrophilic and mesophilic *Chlamydomonas* species"

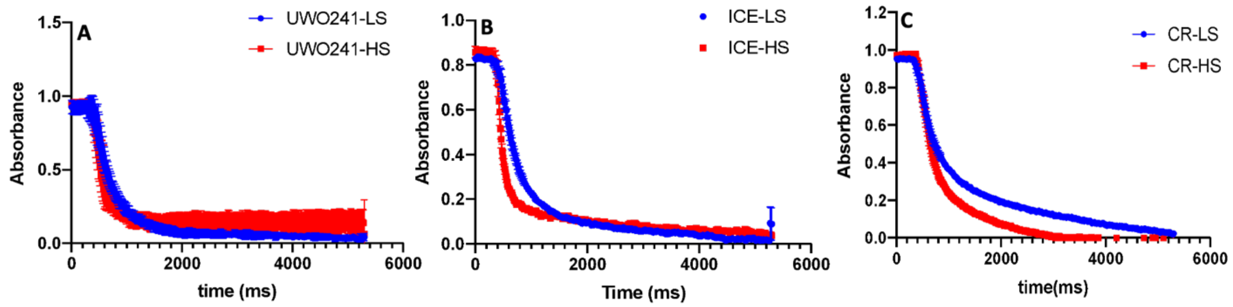


**Supplementary Figure S1: P700 reduction kinetics of the three *Chlamydomonas* species under low and high salinity**. A. UWO241, B. ICE-MDV, C. *C. reinhardtii.* Low salinity (LS) traces are shown in blue. High salinity (HS) traces are shown in red.

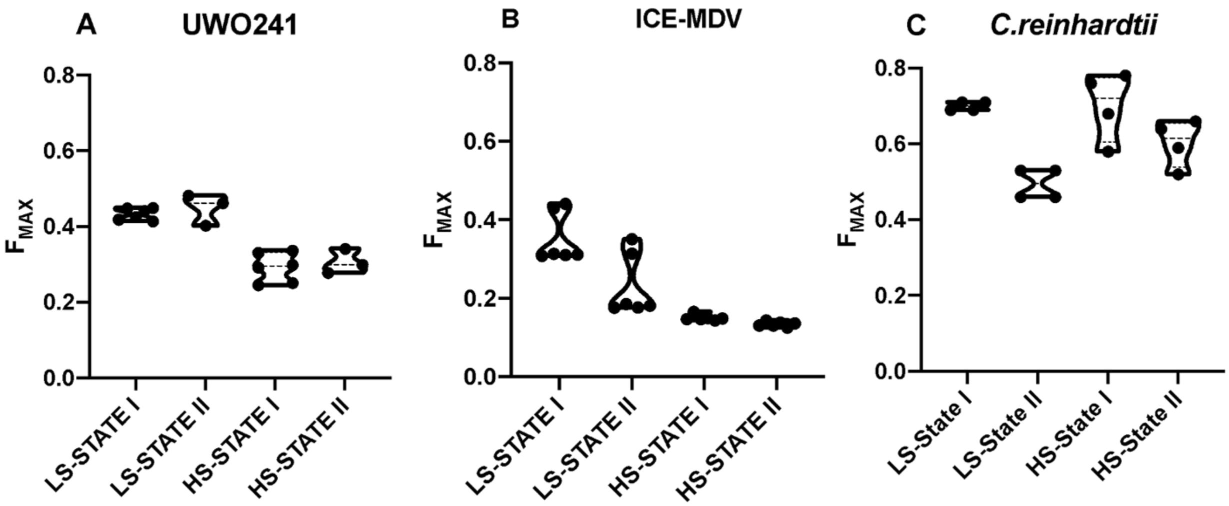


**Supplementary Figure S2**: **PSII state transition test for the three *Chlamydomonas* spp under low (LS) and high (HS) salinity.** The maximum PSII fluorescence values ($F_{MAX}$) are shown for all three strains (UWO241-A, ICE-MDV-B and *C.reinhardtii*-C) under state I and state 2 conditions. State I: DCMU, State II: FCCP. (*n*=4-6, dotted line=mean value)

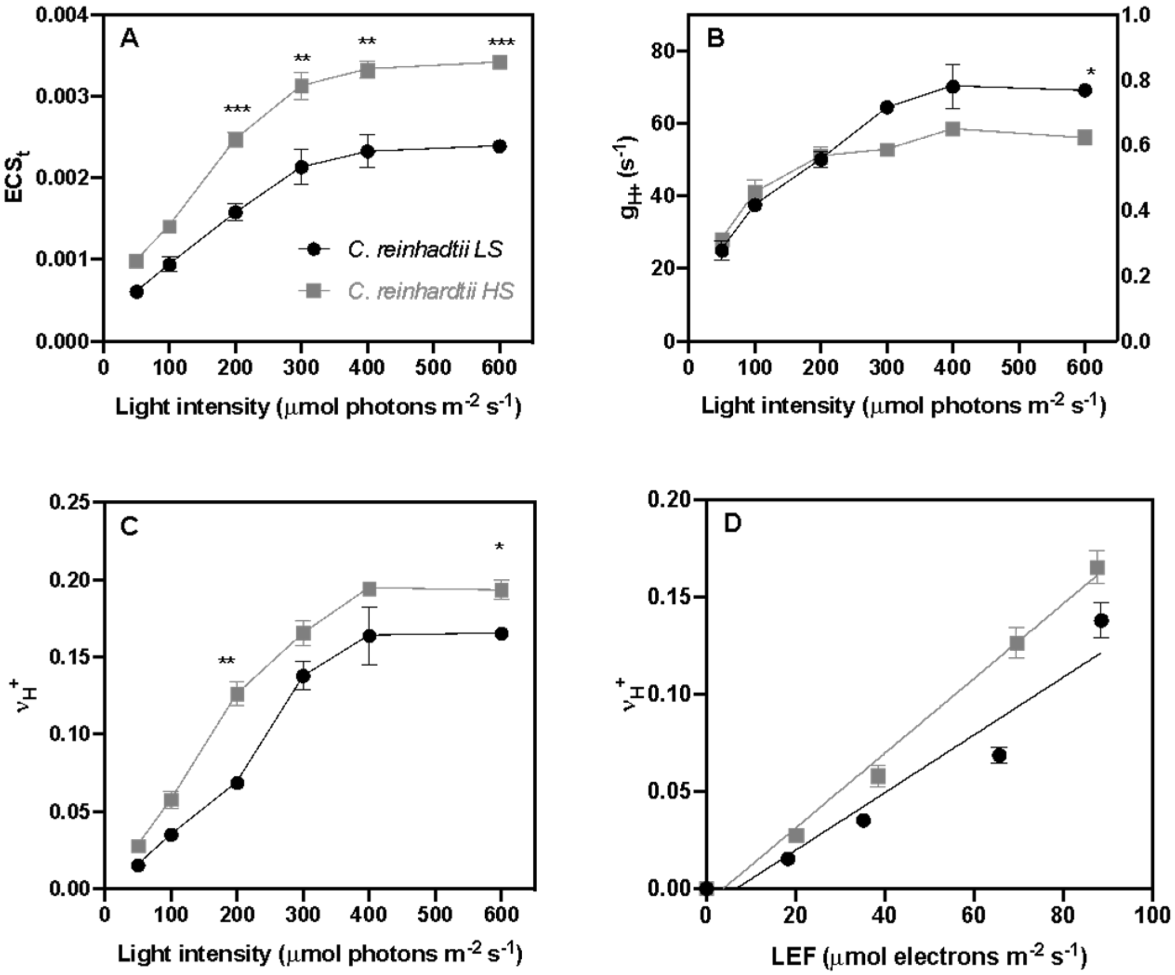


**Supplementary Figure S3: Electrochomic shift (ECS) measurements for *C. reinhardtii* after low and high salinity acclimation.** A. Total proton motive force (pmf) is shown as total change in the ECS signal ($ECS_t$) for low (LS, black) and high (HS, grey) salt acclimated cultures under increasing light intensities. B. ATP synthase conductivity ($g_H^+$) C. Total proton flux ($v_H^+$) and D. Change in total proton flux as a function of linear electron flow (LEF). (*n*=3, SD,* ($p < 0.05$), ** ($p<0.01$), *** ($p<0.005$), **** ($p<0.001$))

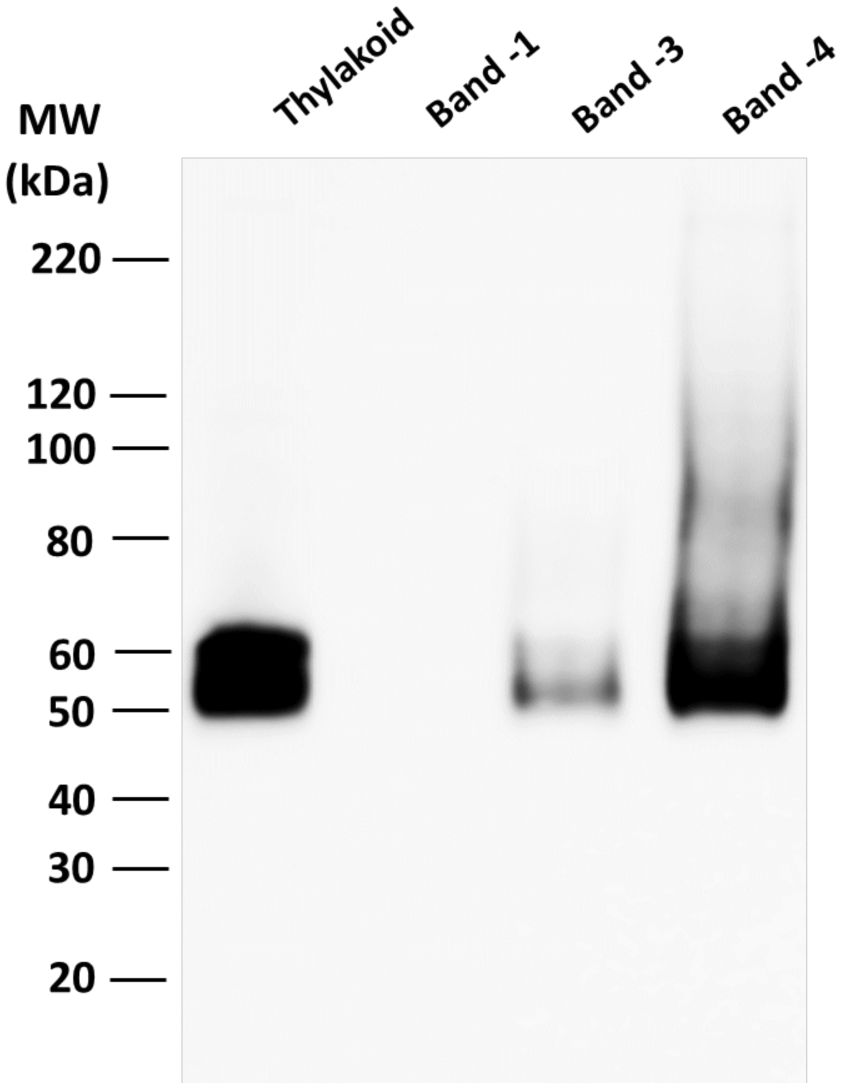


**Supplementary Figure S4: Immunoblot of PsaA in UWO241-HS thylakoids and protein complex fractions collected from sucrose density gradient centrifugation**. Band-1: LHCII complex, Band-3: PSI-LHCI complex, Band-4: Supercomplex. Molecular wt ladder is shown on left.
